# autoStreamTree: Genomic variant data fitted to geospatial networks

**DOI:** 10.1101/2023.05.27.542562

**Authors:** Tyler K. Chafin, Steven M. Mussmann, Marlis R. Douglas, Michael E. Douglas

## Abstract

**Summary:** Landscape genetics is a statistical framework that parses genetic variation within the context of spatial covariates, but current analytical methods typically fail to accommodate the unique topologies and autocorrelations inherent to network-configured habitats (e.g., streams or rivers). autoStreamTree analyzes and visualizes genome-wide variation across dendritic networks (i.e., riverscapes).

**Availability and Implementation:** autoStreamTree is an open source workflow (https://github.com/tkchafin/autostreamtree) that automatically extracts a minimal graph representation of a geospatial network from a provided shapefile, then ‘fits’ the components of genetic variation using a least-squares algorithm. To facilitate downstream population genomic analyses, genomic variation can be represented per-locus, per-SNP, or via microhaplotypes (i.e., phased data). We demonstrate the workflow by quantifying genetic variation in Speckled Dace (*Rhinichthys osculus*) *versus* environmental covariates, with putative adaptive variants subsequently identified.

## 1. INTRODUCTION

Network approaches, particularly those graph-theoretic in nature, are increasingly being used to capture functional ecological or evolutionary processes (e.g., dispersal, gene flow) within/ among habitat patches (Peterson *et al*., 2013). In some cases (e.g., riverscapes) topological patterns are explicitly mirrored by the physical habitat, such that the network structure itself places constraints upon processes such as individual movement (Campbell Grant *et al*., 2007). It is no surprise then, that the importance of network properties such as topological complexity are increasingly implicated as driving evolutionary dynamics in dendritic habitats (Thomaz *et al*., 2016; Chiu *et al*., 2020).

Despite this, quantitative frameworks for modelling the relationships between evolutionary and ecological processes (e.g., through spatio-genetic associations) are predominantly focused on landscapes, and as such often involving mechanistic assumptions which translate poorly to networks. We address this limitation by providing a novel package, autoStreamTree, that facilitates network modeling of genome-scale data. It first computes a graph representation from spatial databases, then analyses individual or population-level genetic data to ‘fit’ distance components at the stream- or reach-level within the spatial network. Doing so within a network context allows the explicit coupling of genetic variation with other network characteristics (e.g., environmental covariates), in turn promoting a downstream statistical process which can be leveraged to understand how those features drive evolutionary processes (e.g., dispersal/ gene flow). We demonstrate the utility of this approach with a case study in a small stream-dwelling fish in western North America.

## 2. PROGRAM DESCRIPTION

### 2.1 Workflow and User Interface

autoStreamTree employs the Python networkx library (Hagberg *et al*., 2008) to parse geospatial input (i.e., large stream networks) into a graph structure with stream segments as *edges*, sampling locations as *endpoints*, and river junctions as *nodes*. Sample data comprise a tab-delimited table of latitude/longitude coordinates, genome-wide variant data in VCF format, and (optionally) a tab-delimited population map. The data structure ‘graph’ on which downstream computations are performed is built as follows: 1) Sample points are ‘snapped’ to nearest river network nodes (i.e., defining endpoints); 2) Shortest paths are identified between each set of endpoints (Dijkstra, 1959); and 3) A minimal network of original geometries, with contiguous *edges* derived by joining individual segments with junctions (*nodes*) retained that fulfill shortest *paths*.

Pairwise genetic distances from VCF-formatted genotypes are derived among individuals, sites, or populations (via *a priori* user-specifications). Options for sequence- and frequency-based statistics are provided (*-d*/*--dist*). Mantel tests are available to quantify correlations among genetic/ hydrologic distance matrices (Mantel, 1967). The primary workflow is a least-squares procedure analogous to that used to compute branch lengths within a neighbor-joining phylogenetic tree(Kalinowski *et al*., 2008). The procedure fits components of the genetic matrix to *k*-segments in a network, such that fitted distance values (*r*) for each segment separating two populations will sum to the observed pairwise matrix value. This provides a distance (*r*_*k*_) for each of *k*-segments as the genetic distance ‘explained’ by that segment.

Workflow steps are controlled through the command-line interface (*-r*/*--run* argument), with results as plain text tables, and plots via the seaborn package(Waskom, 2021). Fitted distances are added to the input geodatabase and exported as .*shp*.

### 2.2 Features

Additional layers of control are provided to minimize pre-processing steps. Users may define individual/ site aggregates: 1) Through a tab-delimited classification file; 2) By automatically deriving group membership geographically; or 3) Using an automated DBSCAN clustering method (Pedregosa *et al*., 2011; Ester *et al*., 1996).

Users may also provide pre-computed genetic distance matrices either directly at individual or locus levels. Built-in options are provided to concatenate single-nucleotide polymorphisms (SNPs) either globally, or per contig/scaffold. Individual-level statistics include uncorrected *p*-distances (i.e., proportion of nucleotide differences/alignment length), and corrected measures [e.g., JC69 (Jukes and Cantor, 1969)], aggregated by site- or at population-level [e.g., as median, arithmetic mean, or adjusted harmonic mean (Rossman, 1990)], or computed as distances via several frequency-based methods [e.g., Jost’s D (Jost, 2008); Da (Nei and Chesser, 1983); Chord distance (Cavalli-Sforza and Edwards, 1967);*θ*_*ST*_ (Weir and Cockerham, 1984)]. autoStreamTree can also be computed per-locus by specifying *-r RUNLOCI*, and with *-c LOC* in the case of phased data to treat linked SNPs to microhaplotypes.

## 3. DEMONSTRATION AND BENCHMARKING

### 3.1 Case Study: Speckled Dace (Rhinichthys osculus)

To demonstrate autoStreamTree, we employed existing SNP data for Speckled Dace (*Rhinichthys osculus*)(Mussmann, 2018; Mussmann *et al*., 2020). Data represent 13,219 SNPs from *N*=762 individuals across 78 localities in the Colorado River ecosystem.

Stream networks were parsed directly as a minimal sub-graph from RiverATLAS, which contains various local-scale environmental/ hydrological features as annotations (i.e., physiography, climate, land-cover, geology, anthropogenic effects) (Linke *et al*., 2019). Genetic distances were computed globally and per-locus among sites as Jost’s *D* (Jost, 2008). To compare with Kalinowski *et al*. (2008), we used unweighted least-squares, iterative negative distance correction, and replicated analyses using linearized *F*_ST_ independently recalculated in Adegenet (Jombart, 2008).

We examined variation in per-locus fitted distances as a function of environmental and anthropogenic covariates, carried over as annotations to RiverATLAS. We reduced *N*=281 hydro-environmental RiverATLAS attributes using forward-selection in adeSpatial (Dray *et al*., 2022), after first removing variables showing significant pairwise correlations. Remaining selected variables were employed in partial-redundancy analysis (pRDA; to ‘partial-out’ hydrologic distance), with outliers as stream segments/SNPs with loadings ±3 standard deviations from the mean.

### 3.2 Results and Comparisons with Other Software

Runtimes are reported for a 2021 Macbook Pro, 16GB memory, 3.2GHz M1 CPU. Time required to calculate/extract a minimal sub-graph containing 118 dissolved *edges* from RiverATLAS (North America shapefile totaling 986,463 original vertices) was 35min. Computing pairwise hydrologic distances required an additional 3sec. Pairwise population genetic distances were computed in ∼24min (linearized *θ*_*ST*_), with Mantel test and distance fitting requiring 11sec and 10sec, respectively. Re-running the entire pipeline per-locus (i.e., *-r RUNLOC*) took 3h 34min. Fitted-*F*_ST_ for autoStreamTree (Fig. 1A) matched that re-calculated using the Kalinowski *et al*. (2008) method (adjusted *R*^2^ = 0.9955; *p* < 2.2*e*-16). However, due to runtime constraints and manual pre-processing for the latter, per-locus distances were not attempted. The pRDA selected four variables, with 221 SNPs (Fig. 1B) and 9 *edges* (Fig. 1C) as outliers.

**Figure 1:**
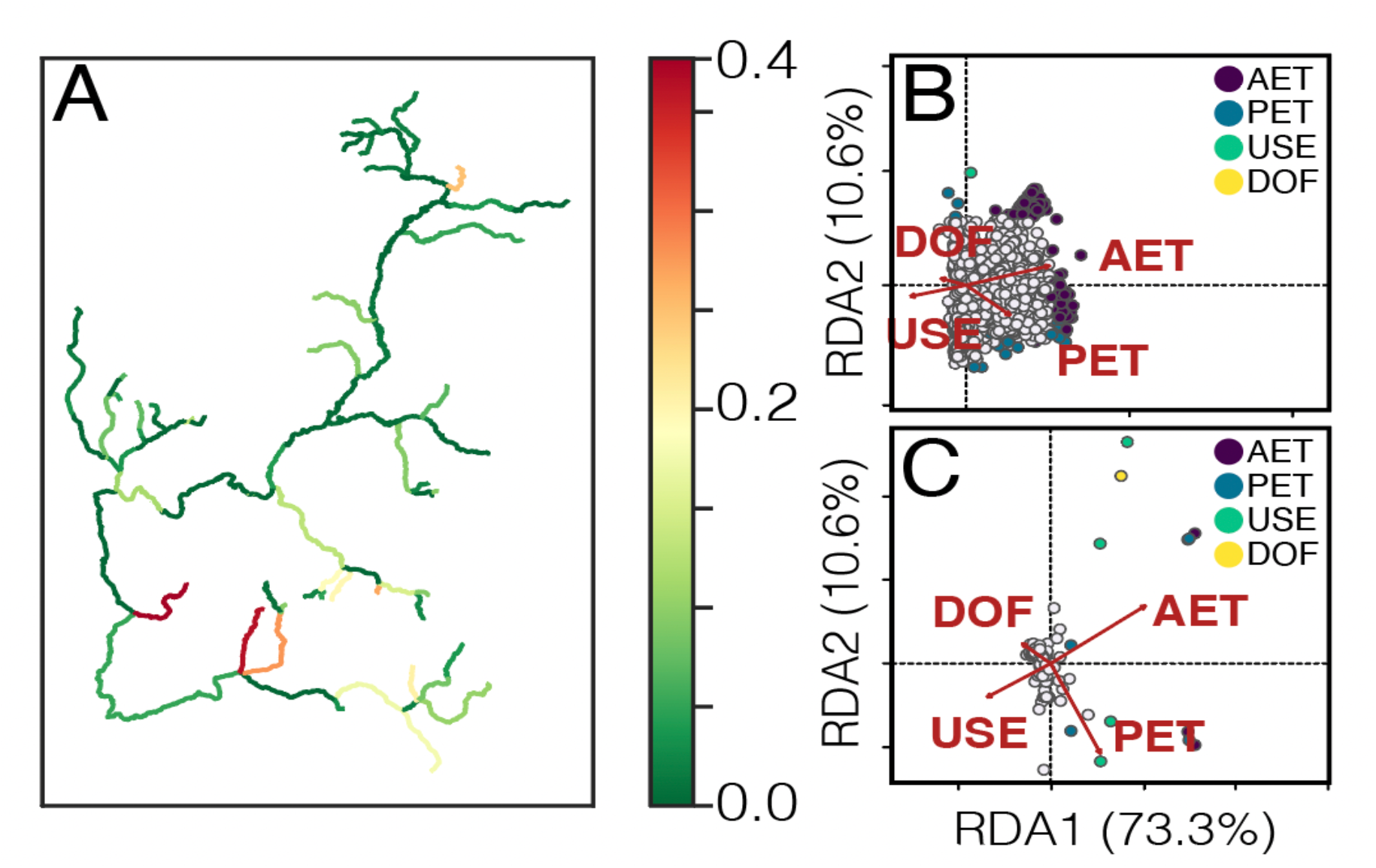
autoStreamTree output. Shown are *F*_ST_ distances fitted onto original stream network (A), variation in per-locus fitted-*F*_ST_ distances via pRDA (controlling for stream length) scaled by loci (B), and by stream segment (C). Outliers highlighted according to most closely correlated explanatory axis and abbreviated as: AET (actual evapotranspiration), PET (predicted evapotranspiration), DOF (degree of fragmentation), and USE (proportion of water for anthropogenic use).

## 4. CONCLUSION

The utility of autoStreamTree was demonstrated with a population genomic dataset as a demonstrative case study. The benefits of the automated approach are underscored by locus-wise microhaplotype *versus* SNP analysis, which in turn feeds into a quantitative framework that allows ‘outlier’ loci exhibiting environmental/ spatial associations within the autocorrelative structure of the network to be detected. This may potentially imply adaptive variation (although not evaluated herein). In addition, the approach is portable to other data types – indeed, any distance matrix that can be appropriately modeled additively can be supplied, and the process is generalizable to any manner of spatial network.

## ACKNOWLEDGEMENTS

Links to non-Service websites do not imply any official U.S. Fish & Wildlife Service endorsement of the opinions or ideas expressed therein or guarantee the validity of the information provided. The findings, conclusions, and opinions expressed in this article represent those of the authors, and do not necessarily represent the views of the U.S. Fish & Wildlife Service.

